# PGBTR: A powerful and general method for inferring bacterial transcriptional regulatory networks

**DOI:** 10.1101/2024.03.08.584073

**Authors:** Wei-Cheng Gu, Bin-Guang Ma

## Abstract

Predicting bacterial transcriptional regulatory networks (TRNs) through computational methods is a core challenge in systems biology, and there is still a long way to go. Here we propose a powerful, general, and stable computational framework called PGBTR, which employs Convolutional Neural Networks (CNN) to predict bacterial transcriptional regulatory relationships from gene expression data and genomic information. PGBTR consists of two main components: the input generation step PDGD and the deep learning model CNNBTR. On the real *Escherichia coli* and *Bacillus subtilis* datasets, PGBTR outperforms other advanced supervised and unsupervised learning methods in terms of AUROC, AUPR, and F1-score. Moreover, PGBTR exhibits greater stability in identifying real transcriptional regulatory interactions compared to existing methods. PGBTR provides a new software tool for bacterial TRNs inference, and its core ideas can be further extended to other molecular network inference tasks and other biological problems using gene expression data.

## Introduction

The growth and differentiation of cells are inseparable from the normal expression of genes, and the gene expression process can be regulated at many stages [1, 2]. Transcriptional regulatory networks (TRNs) refer to a series of molecules and signaling pathways that regulate gene expression at the transcription level [3], defining the regulatory relationships between transcription factors (TFs) and their target genes under different conditions. TRN constitutes a basic framework for understanding the development and physiological responses of an organism, and fully characterized TRNs can predict and explain the dynamic adaptive mechanisms of organisms to environmental or genetic perturbations [3, 4]. In the pre-genomic era, reconstruction of genome-scale TRNs mainly relied on extensive experiments to integrate the one-to-one direct physical interaction of each regulator with its target gene [5]. However, the emergence and rapid development of high-throughput sequencing technology in the post-genomic era has changed the game rules [6]. Massive whole-genome expression data and large-scale TF binding site identification data are constantly increasing. Inferring TRNs from these high-throughput data becomes a long-term core challenge in systems biology [7, 8].

Over the past few decades, many methods have been proposed to reconstruct bacterial TRNs from high-throughput data. These methods can be roughly divided into three categories: unsupervised learning, semi-supervised learning, and supervised learning methods. Among them, unsupervised learning methods occupy a dominant position. Unsupervised learning methods can be further divided into information-based and model-based methods. Information-based methods determine whether there is a regulatory relationship by calculating correlation indicators between gene expressions [9, 10]. Model-based methods build a model that can accurately describe the relationship between genes by fitting gene expression data. These models can be Boolean networks [11, 12], Bayesian networks [13, 14], models based on differential equations [15, 16], logical models [17], and machine learning models [18-20]. In addition to using bulk gene expression data for regulatory inference, more and more unsupervised learning methods are beginning to integrate other types of data to improve method performance or to focus on obtaining more detailed dynamic TRNs [21].

Semi-supervised learning methods often obtain more reliable negative training samples from unlabeled data through inductive and transductive learning, and then use these labeled data for training and prediction through machine learning models. For example, Patel and Wang [22] proposed a semi-supervised method for gene regulatory networks (GRNs) prediction based on two machine learning algorithms: random forest (RF) and support vector machine (SVM). They demonstrated the effectiveness of the method on the regulatory relationships corresponding to four specific transcription factors in *E. coli*.

With continuous research on bacterial TRNs, a substantial number of transcriptional regulatory relationships have been identified, and supervised learning methods have also begun to show their strength in the field of TRN inference. These methods transform the inference of TRN into a binary classification problem, predicting regulatory relationships between TFs and genes. For instance, Mordelet and Vert proposed SIRENE [20], which separately trains an SVM classifier for each TF, and then uses this classifier to predict whether other genes are regulated by the TF. Afterwards, a supervised learning method named GRADIS [23] was designed to reconstruct TRNs using distance distributions obtained from graph representation of transcriptomic data using SVM. GRADIS outperformed basic SVM models and various unsupervised methods on the Dream5 *E. coli* datasets. In recent years, with the continuous development of deep learning, some researchers have begun to use deep learning models to predict bacterial transcriptional regulatory relationships. As the pioneering work of the graph neural network (GNN) model in this field, GRGNN [24] expresses the TRN inference as a graph classification problem. It begins by constructing an initial noisy graph using Pearson correlation coefficients and mutual information derived from gene expression data. Subsequently, the model employs a GNN to train and predict based on these initial subgraphs.

However, these existing methods have limited application scenarios in actual bacterial TRNs inference. For example, for the unsupervised methods, their advantage is that they do not require a priori knowledge of TRNs, making them highly universal. However, the main challenge with unsupervised methods is the interpretation of results. It is difficult to determine a suitable threshold to judge whether there is a regulatory relationship between a TF and a gene. In addition, many studies have also shown that carefully trained supervised models perform better than unsupervised methods [20, 25]. In the field of bacterial TRNs inference, research on supervised learning methods lags far behind that in eukaryotes. The limitations are not only at the data level, including the gaps in available standard datasets and the heterogeneity of expression data sources, but also at the methodological level, including the challenges to effectively utilize various omics data. On the other hand, integrating multiple omics data will bring new problems. The resource-intensive method will be limited by the input data when inferring whole-genome networks or extending to other species. Therefore, developing a powerful and universally applicable computational framework for bacterial TRNs inference remains a great challenge.

Here, we propose a CNN-based method for large-scale bacterial TRNs reconstruction, named PGBTR. It can predict the regulatory relationships between TFs and genes based on gene expression data obtained by microarray or RNA-seq. PGBTR is mainly composed of two parts: Probability Distribution and Graph Distance (PDGD) for input generation and Convolutional Neural Networks for Bacterial Transcriptional Regulation inference (CNNBTR) for model training and prediction. PDGD is a novel and powerful input generation method, which can effectively convert gene expression profiles into input matrices readable by the CNN model. CNNBTR can predict unknown regulatory relationships by learning the complex nonlinear relationships from these matrices. Additionally, CNNBTR also combines the easily available genomic distance to improve model performance.

We comprehensively evaluated PGBTR on the Dream5 datasets (synthetic and *E. coli*) [8], and our newly constructed two bacterial datasets of *Escherichia coli* and *Bacillus subtilis*. Across the three real bacterial datasets, PGBTR is superior to currently available methods, demonstrating its excellent performance in bacterial TRNs inference. Moreover, on the two datasets we constructed, the conservativeness of the positive sample identification results also proves its stability. Overall, our PGBTR method is a powerful, general and stable computational framework for bacterial TRNs inference.

## Materials and Methods

### 2.1 The framework of PGBTR

#### 2.1.1 PDGD matrix

Here we first present a method to transform gene expression data into a matrix form that can be used as input to CNN (**Figure 1A**), Probability Distribution and Graph Distance (PDGD). We begin by selecting a gene pair (gene A encoding a transcription factor and another gene b) and convert their expression data into a 32 × 32 two-dimensional histogram. In this histogram, the abscissa is the expression level of gene A, and the ordinate is the expression level of gene b. Next, the ranking of their gene expression data within the entire gene expression profile is then transformed into a second 32 × 32 two-dimensional histogram. This ranking-based feature is used because it can reduce systematic bias in transcript counting [26]. Then the *K*-means algorithm is used to cluster the entire gene expression profile into 50 categories, and the cluster centroids of gene A and gene b are extracted and converted into 50 points 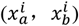 (*i* = 1, 2, 3, …, 50) in the two-dimensional coordinate system. We calculate the Euclidean distance between these points by **Eq. 1**:

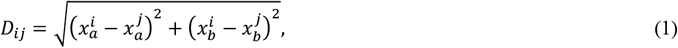

where *i* = 1, 2, 3, …, 50 and *i* < *j* ⩽ 50. Subsequently, fill the first 1024 ones of the calculation results into a third 32 × 32 two-dimensional matrix, and finally splice these three matrices into a 32 × 32 × 3 three-dimensional matrix, which is the PDGD matrix. A suitable matrix size is very important for subsequent prediction tasks. Smaller matrices will lead to the loss of data features, while larger matrices will introduce additional noise, so we compared the model performance under different matrix sizes and finally determined the size of the PDGD matrix to be 32 × 32 × 3.

**Figure 1.**
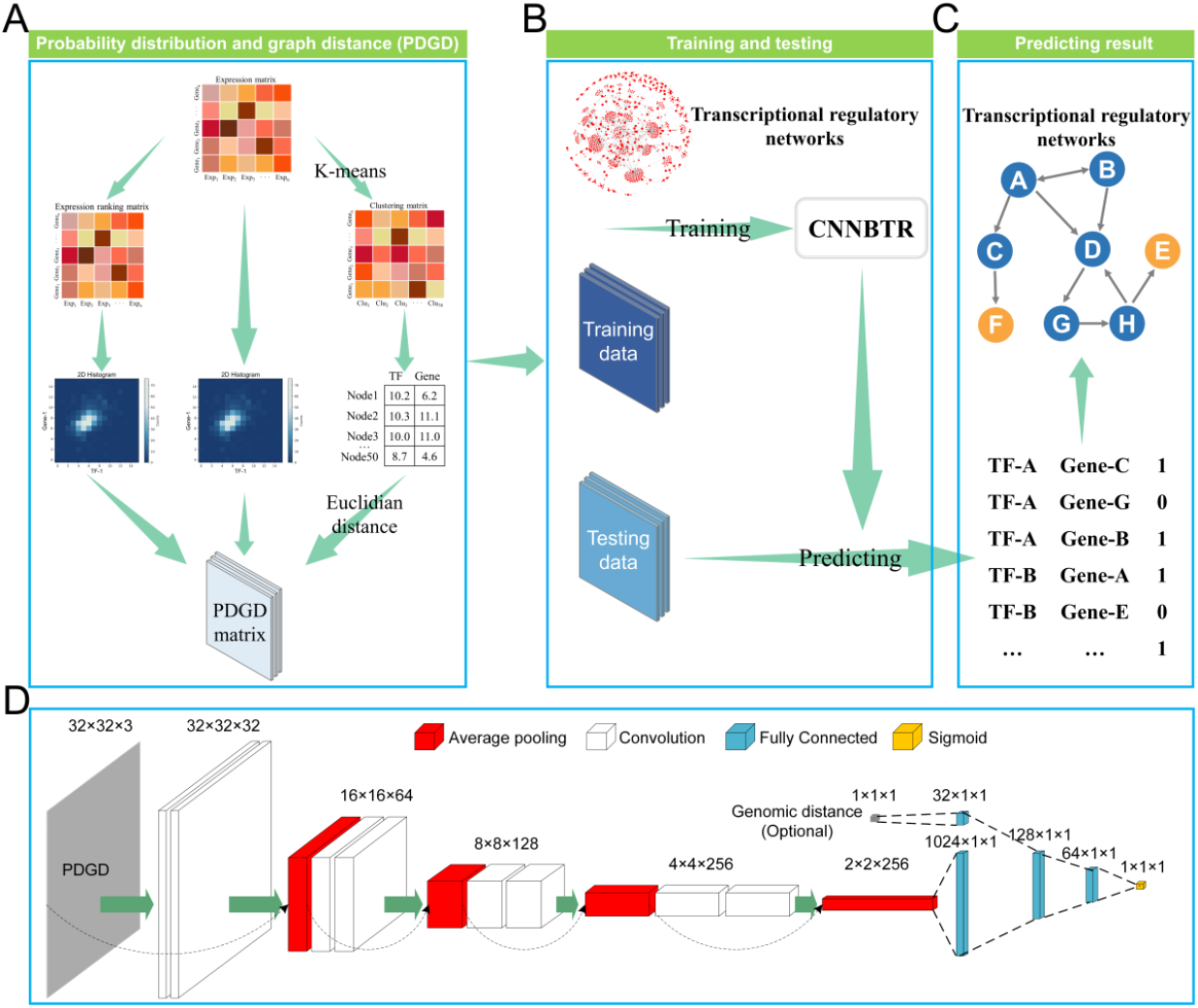
The workflow of PGBTR. (A) Convert gene expression data (by microarray or RNA-seq) into PDGD (probability distribution and graph distance) matrices. (B) Divide the PDGD matrices into training and testing datasets. (C) Obtain the predicted labels (0 or 1) of the TF-gene pairs. Based on these labels, TRN can be constructed. (D) The neural network structure of CNNBTR (convolutional neural networks for bacterial transcriptional regulation inference).

#### 2.1.2 The architecture of CNNBTR

CNNBTR (convolutional neural networks for bacterial transcriptional networks inference) consists of three layers: input layer, hidden layer, and output layer (**Figure 1D**). The input layer is the above-mentioned 32 × 32 × 3 PDGD matrix, and the hidden layer is composed of four ResNet modules and two fully connected layers. The ResNet module refers to the classic ResNet [27], which includes a convolutional layer using a 3 × 3 convolutional kernel (**Eq. 2**) and an average pooling layer using a 2 × 2 pooling kernel (**Eq. 3**). We use the rectified linear activation function (ReLU) (**Eq. 4**) as the activation function across the entire hidden layer, while the final output layer uses Sigmoid function (**Eq. 5**) as the activation function for binary classification. The function of the convolutional kernel is defined as follows:

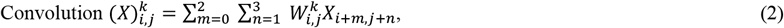

where *X* is the input from the previous layer, (*i, j*) is the output position, *k* is the convolutional filter index, and *W* is the filter matrix of size 3 × 3.

The average pooling layer uses the following function:

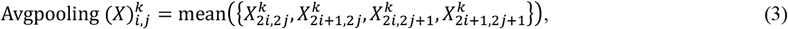

where *X* is the input of the previous layer, (*i, j*) is the output position, and *k* is the index of the pooling kernel. The ReLU function is defined as follows:

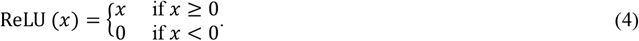

The Sigmoid function is defined as follows:

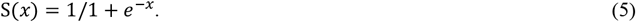

### 2.2 Benchmark datasets

In this paper, we selected the *E. coli* and simulation datasets from the Dream5 challenge as the experimental datasets. In addition, we also constructed new standard datasets for *E. coli* and *B. subtilis* based on the latest research. **Table 1** shows the details of the datasets used in this study.

**Table 1.** Information on the datasets used in this study.

| Expression data | Gold standard TRN |
| --- | --- |

|  | Genes | Samples | TFs | Links |
| --- | --- | --- | --- | --- |
| Dream5_Silico | 1643 | 805 | 195 | 4012 |
| Dream5_Ecoli | 4511 | 805 | 334 | 2066 |
| RegulonDB_Ecoli | 4270 | 1035 | 220 | 5316 |
| Subtiwiki_Bsubtilis | 4325 | 265 | 185 | 3033 |

#### 2.2.1 Dream5 challenge datasets

This study first evaluated all methods on the *E. coli* dataset and simulated dataset from the Dream 5 challenge [8]. Both datasets include a microarray gene expression profile, a gold standard TRN, a list of transcription factors, and a list of gene id transitions. Among them, the gold standard network of the simulated dataset contains 4012 positive samples (regulatory pairs) and 274,380 negative samples (non-regulatory pairs), and the gold standard network of the *E. coli* dataset contains 2066 positive samples and 150,214 negative samples.

#### 2.2.2 Self-constructed *E. coli* dataset — RegulonDB_Ecoli

In recent years, the RNA-seq data for *E. coli* under various experimental conditions have been generated and many new transcriptional regulatory relationships in this species have been discovered. We used the TF-gene transcriptional regulatory relationships recorded in RegulonDB 11.0 [28] to construct a new gold standard transcriptional regulatory network. Gene expression data were derived from the precise1K dataset [29], a dataset of 1035 *E. coli* RNA-seq samples that includes a wide range of genetic perturbations and environmental conditions. In order to unify the gene information in the TRN and bulk RNA-seq data, we performed the following preprocessing steps: (1) remove the self-regulatory relationships in the TRN; (2) remove the regulatory relationships without gene expression data; (3) treat TFs encoded by multiple genes as a collective entity, with their expression level being the average of the constituent gene expression levels. Finally, the TRN in the RegulonDB_Ecoli dataset we constructed contains 5316 positive samples.

#### 2.2.3 Self-constructed *B. subtilis* dataset — Subtiwiki_Bsubtilis

To verify the applicability of the model in other species, we constructed an additional dataset using *B. subtilis*, which is a well-studied Gram-positive bacterium. We downloaded the gold standard TRN of *B. subtilis* from Subtiwiki 4.0 [30]. In addition, we obtained the RNA-seq dataset for *B. subtilis* from the study of Rychel et al. [31]. We applied the same preprocessing steps as those used for the RegulonDB_Ecoli dataset, and the final TRN contains 3033 positive samples.

### 2.3 Evaluated methods

We compared PGBTR with other published TRN inference methods, including two supervised learning methods and three unsupervised learning methods.

#### 2.3.1 Supervised learning methods

GRADIS [23] is an advanced TRN inference method based on SVM. GRADIS first uses the *K*-means algorithm to reduce the dimensionality of gene expression data, then extracts features from the gene expression data through graph embedding. Finally, the extracted features are input into an SVM model to perform binary classification on regulatory interactions. GRGNN [24] is an advanced TRN inference method based on GNN. GRGNN formulates TRN inference as a graph classification problem, and predicts the regulatory relationships of TF-gene pairs under a supervised or semi-supervised framework.

#### 2.3.2 Unsupervised learning methods

Because there are few supervised learning methods that can be used for bacterial TRN inference, in addition to supervised learning methods, we also selected three unsupervised learning methods that performed well on the Dream5_Ecoli dataset for comparison, including AGRN [32], GENIE3 [19] and PROTIA [33]. AGRN is a gene regulatory network inference method based on integrated machine learning, which uses the Shapley value to give regulatory scores between gene pairs. GENIE3 is a widely used method for gene regulatory network inference based on RF. The strength of PROTIA which based on robust precision matrix estimation lies in its fast execution speed and excellent performance [33].

### 2.4 Training and testing strategy

For supervised learning methods, the datasets from Dream5 challenge provided a large number of negative samples, while for the two datasets of RegulonDB_Ecoli and Subtiwiki_Bsubtilis constructed in this study, a negative sample pool is generated using gene pairs composed of TFs and the genes other than their target genes. Due to the potential impact of imbalanced datasets on supervised learning models, we randomly sample ten times from the negative sample pool to obtain the same number of non-regulatory pairs (negative samples) as the number of known regulatory relationships (positive samples). In the end, we constructed ten balanced datasets for each bacterium.

In the field of computational biology, especially for tasks such as TF-gene pairs classification, the use of supervised learning methods requires special attention to data leakage, because the data overlap between the training and testing datasets will cause the performance of the model on the testing set to be overestimated [34]. When evaluating TRN inference methods, there are two widely used dataset partitioning strategies. The first is to perform cross-validation on the entire dataset indiscriminately. The second is to divide the entire dataset into three parts according to different TFs, and each part of the dataset contains similar number of TFs. Then, three-fold cross-validation is performed, with two parts as training set and one part as testing set. In practical TRN inference, it is common that not all the target genes of certain TFs have been discovered. Thus, compared with the second strategy, the first strategy is closer to practical application. Therefore, the first dataset partitioning strategy is used (**Figure 1B**), and five-fold cross-validation on each balanced dataset is conducted to evaluate the performance of the model, namely, each benchmark dataset is randomly divided into a training set and a testing set according to an 8: 2 ratio, and for each training set, 20% of the training data was randomly selected as a validation set.

For unsupervised learning methods, the obtained results typically consist of regulatory possibility scores for all possible TF-gene pairs. In order to make a fair comparison with the supervised learning methods, we did not calculate various metrics for the top 10,000 regulatory pairs as in the Dream5 challenge. Instead, the corresponding TF-gene pairs of each testing set in supervised learning methods are extracted from all the results to calculate various evaluation metrics.

### 2.5 Metrics for evaluation

To comprehensively assess the performance of all methods, we utilized three key metrics: area under the ROC curve (AUROC), area under the Precision-Recall curve (AUPRC) and F1-score. In binary classification tasks, AUROC is one of the most commonly used metrics to measure model performance. Besides, considered that in TRNs inference tasks, researchers are more concerned with the prediction results of positive samples, and the reliability of negative sample data is much lower than that of positive samples. Therefore, we also considered AUPRC and F1-score, two evaluation indicators that focus on positive samples. In the supervised learning methods, we chose the median of the range of predicted values, and for the unsupervised learning methods, we chose the median of the gene pair scores in the testing set as the threshold for calculating F1-score.

## Results

### 3.1 Implementation of PGBTR

To predict bacterial TRNs, we developed PGBTR, a general computational framework for predicting whether regulatory relationships exist between TFs and genes (**Figure 1**). PGBTR includes two main steps: (1) firstly, the expression data of each gene pair (gene A encoding a TF and another gene b) in the dataset are converted into a PDGD matrix; (2) then, input the generated PDGD matrices into a deep learning model — CNNBTR for training and prediction.

### 3.2 Effectiveness of PDGD

For the application of deep learning models in TRN inference, it is usually necessary to convert raw data into a form that matches the model, a process called input generation. The input generation method will directly affect the quality of the input data, thereby significantly impacting model performance. Therefore, we compare the performance of the NEPDF (Normalized Empirical Probability Distribution Function) method [35] and our PDGD method on our CNNBTR model. Firstly, we use the NEPDF and PDGD methods to convert the gene expression data in the RegulonDB_Ecoli and Subtiwiki_Bsubtilis datasets into matrices of different sizes, and then input these matrices into the CNNBTR model for training and prediction. As shown in **Figure 2**, on both datasets, the PDGD method performs better on the CNNBTR model than the NEPDF method, regardless of matrix sizes. In addition, across the three sizes of matrices: 8 × 8, 16 × 16 and 32 × 32, the performance of the model improves as the matrices size increases. Due to the limited number of bacterial gene expression samples, we did not consider larger-sized matrices, such as 64 × 64. We hope that as the bacterial gene expression data continue to increase, we can try larger-scale input matrices in the future. Our PDGD method provides a novel approach for input generation of gene expression data for deep learning models, especially CNN models, which is expected to help TRN inference and other biological problems that need using gene expression data.

**Figure 2.**
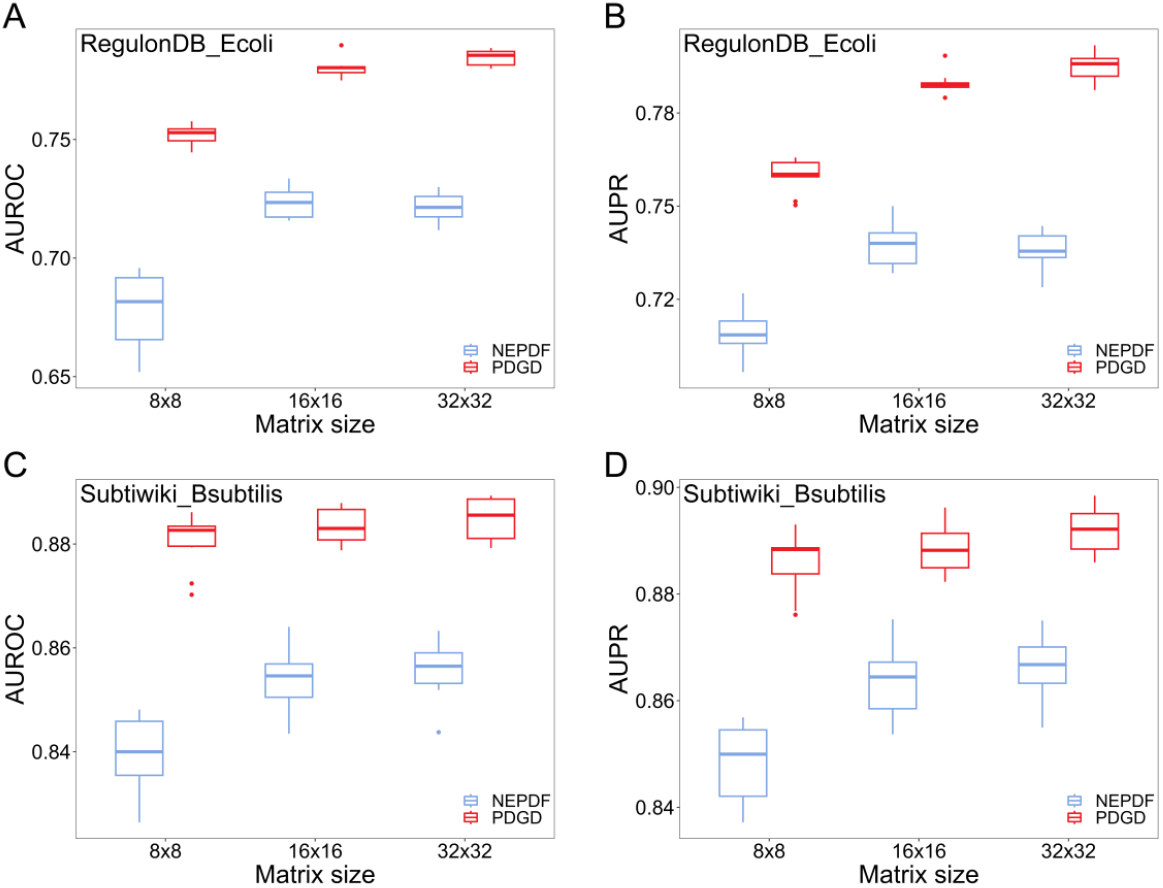
Performance of the two input generation methods (PDGD and NEPDF) on CNNBTR. (A, B) AUROC and AUPR comparison of PDGD (red) and NEPDF (blue) on the RegulonDB_Ecoli dataset. (C, D) AUROC and AUPR comparison of PDGD (red) and NEPDF (blue) on the Subtiwiki_Bsubtilis dataset.

### 3.3 PGBTR performs well in TRN prediction

To evaluate the performance of PGBTR in bacterial TRN inference, we ran PGBTR and five other evaluated methods on the Dream5 datasets, RegulonDB_Ecoli and Subtiwiki_Bsubtilis datasets (see **Materials and Methods** for details). Since all genes in the Dream5 datasets are anonymous, the genomic distance (the difference in base pair of the positions of two genes on the linear genome) information between TF-gene pairs cannot be used, we adopted a reduced version of the PGBTR model on the Dream5 datasets (the PGBTR_dream model which does not need this information).

#### 3.3.1 On the Dream5 datasets

**Table 2** shows the performance of all the evaluated methods on the Dream5 datasets. This table presents the average and standard deviation of the five-fold cross-validation results based on the ten balanced datasets.

**Table 2.**
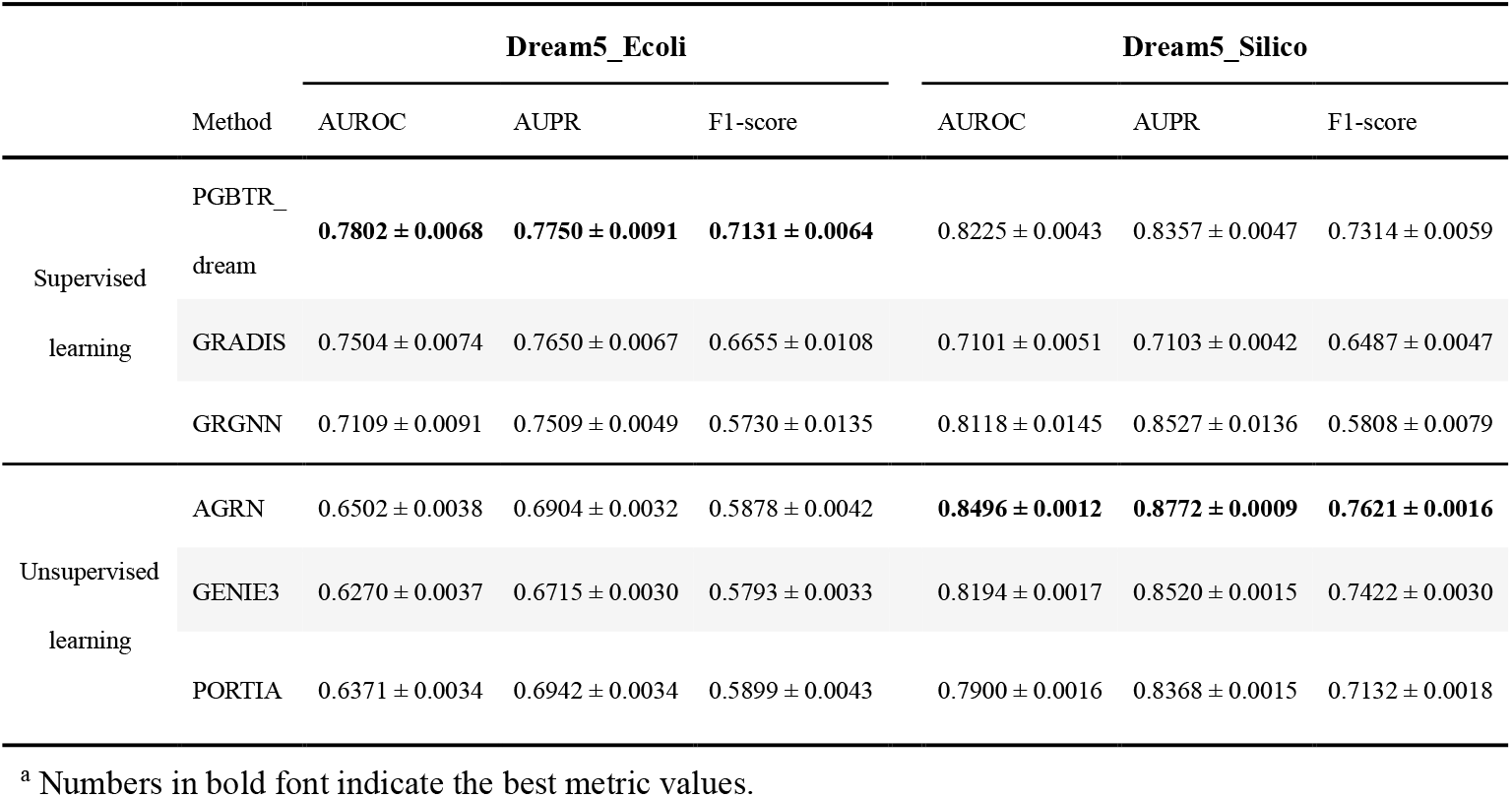
Performance of each method on the Dream5 challenge datasets^a^.

On the Dream5_Ecoli dataset, the PGBTR_dream model achieves the best performance in all the three metrics: AUROC, AUPR, and F1-score. Meanwhile, consistent with previous research [20, 25], supervised learning methods overall outperform unsupervised learning methods on the Dream5_Ecoli dataset.

On the Dream5_Silico dataset, the AUROC of the PGBTR_dream model ranks second among all methods, superior to the other two supervised learning methods. Notably, it is interesting that the unsupervised methods perform better than supervised learning methods overall, and AGRN achieves first place in all three metrics. This does not conflict with the findings of Maetschke et al. [25], who compared the performance of supervised and unsupervised learning methods on small-scale networks (30 genes), while our comparison was made on synthetic datasets of large-scale networks (1643 genes). For such results, we suppose that the relationship of expression levels between TFs and target genes in the synthetic dataset is much simpler than that in the real data, making it easier to directly distinguish whether there is a regulatory relationship. Additionally, input generation steps introduce new noises while transforming the expression data into specific input formats. As a result, the performance of the trained models is worse than that of the unsupervised methods that directly infer regulatory relationships from expression data without relying on labels.

#### 3.3.2 On the RegulonDB_Ecoli and Subtiwiki_Bsubtilis datasets constructed in this study

While the Dream5 datasets are widely used for the evaluation of TRN inference methods, they are already datasets from twelve years ago. Therefore, in order to more comprehensively evaluate the PGBTR model in predicting bacterial TRNs, we selected the two most representative bacteria, *E. coli* (Gram-negative) and *B. subtilis* (Gram-positive), and constructed the RegulonDB_Ecoli and Subtiwiki_Bsubtilis datasets using relevant data from public databases. **Table 3** presents the performance of all the methods on these two datasets. Likewise, this table displays the average and standard deviation of five-fold cross-validation results on all the ten balanced datasets. Unlike the Dream 5 datasets where all gene names are hidden, all the gene names in real bacterial TRNs prediction tasks are publicly visible, which provides convenience for our model to incorporate more priori information. We tested both PGBTR that incorporates genomic distance information and the PGBTR_dream model that does not incorporate this information. It can be clearly seen that PGBTR achieves the best performance in all three metrics on these two datasets. Both AUROC and AUPR are at least 3% higher than the second-ranking method, while the F1-score is approximately 6% higher. Moreover, incorporating genomic distance information led to a certain degree of performance improvement, indicating that our PGBTR method possesses strong generalization capability and significant potential.

**Table 3.** Performance of each method on the RegulonDB_Ecoli and Subtiwiki_Bsubtilis datasets^a^.

|  | Method | RegulonDB_Ecoli |  |  | Subtiwiki_Bsubtilis |  |  |
| --- | --- | --- | --- | --- | --- | --- | --- |
|  |  | AUROC | AUPR | F1-score | AUROC | AUPR | F1-score |
| Supervised learning | PGBTR | **0.7847 ± 0.0032** | **0.7948 ± 0.0044** | **0.7178 ± 0.0042** | **0.8849 ± 0.0037** | **0.8920 ± 0.0040** | **0.8143 ± 0.0039** |
| | PGBTR<br>_dream | 0.7716 ± 0.0052 | 0.7762 ± 0.0052 | 0.7038 ± 0.0055 | 0.8708 ± 0.0043 | 0.8764 ± 0.0046 | 0.7976 ± 0.0034 |
| | GRADIS | 0.7429 ± 0.0062 | 0.7509 ± 0.0069 | 0.6513 ± 0.0068 | 0.8456 ± 0.0019 | 0.8523 ± 0.0041 | 0.7513 ± 0.0035 |
| | GRGNN | 0.7370 ± 0.0121 | 0.7673 ± 0.0104 | 0.5925 ± 0.0096 | 0.8122 ± 0.0120 | 0.8292 ± 0.0102 | 0.6847 ± 0.0101 |
| Unsupervised learning | AGRN | 0.6030 ± 0.0032 | 0.6475 ± 0.0041 | 0.5689 ± 0.0026 | 0.6257 ± 0.0031 | 0.6616 ± 0.0032 | 0.5779 ± 0.0035 |
| | GENIE3 | 0.6006 ± 0.0033 | 0.6425 ± 0.0045 | 0.5650 ± 0.0030 | 0.6025 ± 0.0053 | 0.6373 ± 0.0043 | 0.5588 ± 0.0042 |
| | PROTIA | 0.5989 ± 0.0030 | 0.6386 ± 0.0034 | 0.5632 ± 0.0027 | 0.5947 ± 0.0023 | 0.6569 ± 0.0024 | 0.5599 ± 0.0025 |

<sup>a</sup> Numbers in bold font indicate the best metric values.

### 3.4 PGBTR is more stable in predicting transcriptional regulatory pairs

Due to the highly imbalanced nature of real bacterial TRNs inference, constructing a balanced dataset through random negative sampling will make the model evaluation more scientifically reasonable, but what we are more concerned about is the accuracy and stability of positive sample predictions. Therefore, we counted the number of correct predictions of true transcriptional regulatory interactions (positive samples) in all balanced datasets for the three supervised learning methods. As shown in **Figure 3**, PGBTR outperforms the other two supervised learning methods by correctly predicting transcriptional regulatory interactions at least eight, nine, and even ten times on both the RegulonDB_Ecoli and Subtiwiki_Bsubtilis datasets. This result indicates that PGBTR is more stable in the recognition of true transcriptional regulatory interactions.

**Figure 3.**
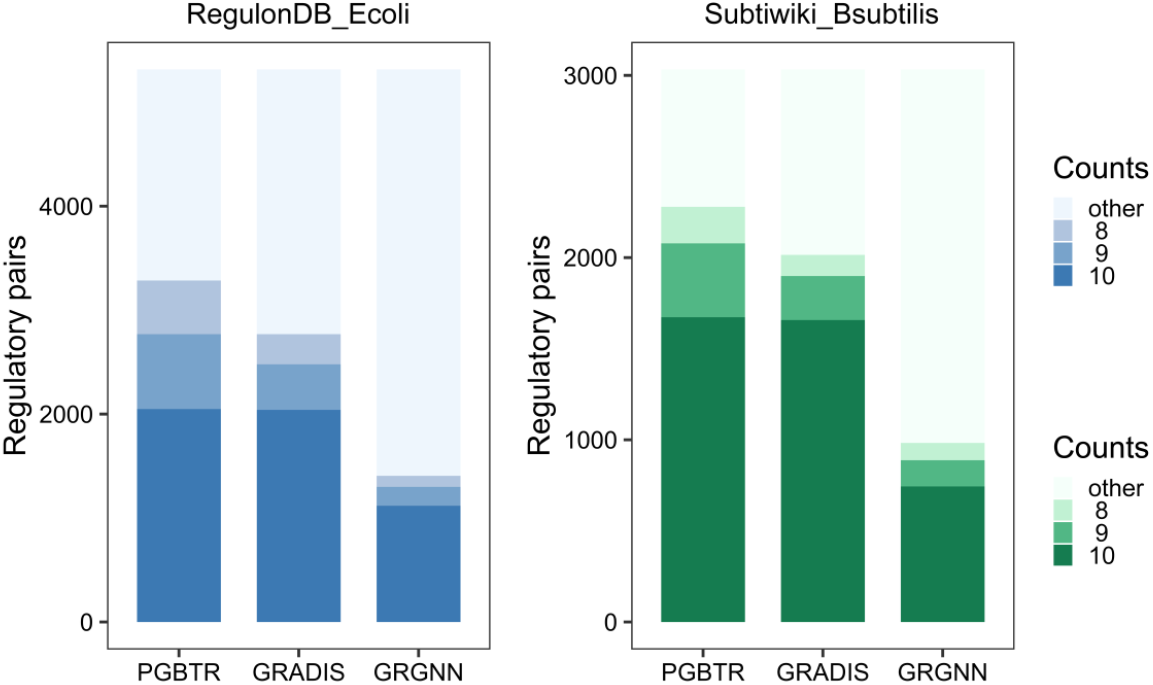
Comparison for the stability of positive sample identification among three supervised learning methods (PGBTR, GRADIS, GRGNN). On the RegulonDB_Ecoli and Subtiwiki_Bsubtilis datasets, the number of times that three supervised learning methods correctly predict for all transcriptional regulatory pairs (positive samples) in ten different balanced datasets constructed. The number of regulatory pairs that are predicted correctly at least 8 times out of ten different balanced datasets is the sum of the number of regulatory pairs that are predicted correctly 8 times, 9 times, and 10 times. The larger this value is, the more stable the model is in predicting positive samples.

## Discussion

With continued efforts, new regulatory relationships are constantly being discovered, and more and more computational methods are developed for the inference of TRNs. However, our understanding on TRNs is still only the tip of the iceberg, even in prokaryotes, which are considered to have simpler TRNs. For example, for the most well-studied bacteria, *E coli*, it is estimated that only 10% to 30% of all regulatory relationships are currently known [36]. Moreover, considering that computational methods for bacterial TRNs inference are far inferior to those in eukaryotes, there is a continued need for the development of advanced computational approaches to better predict bacterial TRNs.

To this end, we propose PGBTR, a general CNN-based computational framework for bacterial TRNs inference. On one hand, the innovation of PGBTR lies in its novel input generation method PDGD, which can preserve the characteristics of the original data as much as possible. One the other hand, PGBTR uses the CNNBTR model inspired by ResNet [27] as the prediction model, incorporating additional connection layers that leverage readily available genomic information. PGBTR stands out for its broad applicability, excellent performance, and considerable potential for scalability.

We conducted a comprehensive comparison of PGBTR with two existing supervised learning methods and three widely-used unsupervised learning methods on four distinct datasets. The results on the Dream5 synthetic dataset indicate that the regulatory relationships in synthetic data differ from those in real large-scale bacterial TRNs, and supervised learning methods that rely on input generation will perform slightly worse than unsupervised learning methods that infer relationships directly from expression data.

Furthermore, PGBTR demonstrates impressive performance on the other three datasets, showcasing its powerfulness, and stability in predicting real bacterial transcriptional regulatory relationships. Firstly, PGBTR has achieved the best performance in all the three metrics on the three datasets of Dream5_Ecoli, RegulonDB_Ecoli, and Subtiwiki_Bsubtilis. On the RegulonDB_Ecoli and Subtiwiki_Bsubtilis datasets, both AUROC and AUPR are at least 3% higher than the other methods, while F1-score is at least 6% higher than the other methods. Secondly, for different balanced datasets constructed by random negative sampling, PGBTR exhibits superior stability in identifying true transcriptional regulatory relationships compared to the other two supervised learning methods. Finally, the performance of PGBTR can be improved to a certain extent by combining genomic distance information, which also proves its profound development potential.

Certainly, there are still several limitations that can be improved when PGBTR is practically used in bacterial TRNs inference. On the one hand, since PGBTR is essentially a supervised learning method, it cannot be generalized to bacteria with few known regulatory relationships. To overcome this limitation, we plan to explore transfer learning and meta-learning approaches. On the other hand, PGBTR mainly identifies regulatory relationships from bulk gene expression data, but in fact these data are not enough to predict the complex and dynamic regulatory relationships between TFs and their target genes in bacteria. In the future, we aim to integrate additional omics data to expand and enhance the function of PGBTR.

## Availability

The PGBTR software is available at https://github.com/mbglab/PGBTR

## Acknowledgements

This work was supported by the National Natural Science Foundation of China (Grant 31971184).

